# Excretory-secretory products from the parasite *Schistocephalus solidus* enhance viability and function of threespine stickleback splenocytes

**DOI:** 10.1101/2025.06.16.659396

**Authors:** Alexandra Collias, Annie Cary, Natalie C Steinel

**Affiliations:** Department of Biological Sciences, University of Massachusetts Lowell, Lowell, MA, USA; Center for Pathogen Research and Training, University of Massachusetts Lowell, Lowell, MA, USA

## Abstract

Helminths can modulate the immune systems of their hosts through the release of soluble molecules known as excretory-secretory products (ESPs). To better understand the extent and mechanisms behind helminth-mediated immunosuppression, we used an emerging model of host-parasite interactions, threespine stickleback, *Gasterosteus aculeatus*, and its tapeworm parasite *Schistocephalus solidus*. We examined the impacts of exposure to ESPs of *S. solidus* (SsESPs) originating from four Canadian lakes on the viability and function of threespine stickleback splenocytes. We found that 24 hours of exposure to low concentrations of SsESPs significantly increased overall splenocyte viability with the greatest impact on the lymphocyte population. While SsESP exposure did not alter baseline splenocyte respiratory burst activity, SsESP-treated splenic cultures demonstrated significant increases in ROS production in response to phorbol 12-myristate 13-acetate (PMA) stimulation, suggesting that SsESPs may lower the threshold for activation of the respiratory burst. These results in splenocytes contrast with previous studies demonstrating that SsESPs suppress head kidney leukocytes (HKL) viability and function, suggest that *S. solidus*-derived excretory secretory products may have cell-or tissue-specific immunomodulatory effects, and highlight the importance of studying host-parasite interactions across diverse immune tissues.

## 1. Introduction

Parasites have evolved evasion strategies to dampen host immunity to promote their own survival, however these changes can compromise the host, leading to negative health outcomes including co-infections and elevated mortality rates^1–3^. On the other hand, the reduction of host immunity by helminths can have immunoregulatory effects, alleviating the symptoms of autoimmune diseases, allergy, and inflammation^4^. Parasitic helminths exert systemic influence on their host’s immune system through the production of soluble mediators known as excretory-secretory products (ESPs)^5–7^. ESP composition and function vary with parasite species, but are generally comprised of proteins, lipids, and extracellular vesicles (EVs) which can contain other small molecules such as microRNAs^6–11^. Fish helminth ESPs can suppress both innate and adaptive immunity through diverse mechanisms, including mimicking host cytokines^12,13^, reducing leukocyte recruitment^14^ and/or proliferation^15^, and down-regulating immune gene expression^16,17^. While there is evidence that fish helminth ESPs can alter immunity of their fish hosts, these studies have been limited to a handful of species.

To broaden our understanding of ESP-mediated immunomodulation, the present study examines the effect of *Schistocephalus solidus* ESPs (SsESPs) on its host, threespine stickleback. *S. solidus* is a trophically transmitted cestode with a complex life cycle involving three hosts: a cyclopoid copepod, threespine stickleback (obligate intermediate host), and a piscivorous bird (terminal host)^18^. Within 24-48 hours of ingesting an infected copepod, the *S. solidus* proceroid penetrates the stickleback’s gastrointestinal tract and establishes infection in the body cavity^18^. *S. solidus* infection can downregulate stickleback immune gene expression^19,20^ and facilitate co-infection with other parasites^21^, suggesting that this cestode has immunosuppressive effects. Recent analysis of the *S. solidus* secretome detected several proteins that are suspected to play important roles in host-parasite interactions, including the suppression of host immunity^22,23^. However, to date, only one study has investigated the role of SsESPs in host immune modulation. Scharsack et al demonstrated the effects of conditioned media from cultured European *S. solidus* on head kidney cultures, finding a reduction in lymphocyte viability and leukocyte respiratory burst^24^. While this study suggests that SsESPs may dampen immunity in infected stickleback, it was limited to a single immune tissue, the head kidney, a hematopoietic organ primarily composed of hematopoietic cells, neutrophils, and B cells^25^. However, immune cells are found in many sites in fish (spleen, thymus, nasopharynx, gills, axilla, bursa)^26^ and the extent of SsESP-mediated modulation on these other tissues and immune populations are unknown.

Therefore, to broaden our understanding of *S. solidus*-mediated immune modulation and potential mechanisms of parasite-mediated immune manipulation, we assessed the effects of North American SsESPs on stickleback splenocyte viability and function *in vitro*. We found that SsESPs from all locations enhanced stickleback splenocyte viability compared to splenocytes cultured in media alone, an effect detectable even at low concentrations. Immune cell subset analysis showed that SsESPs had the greatest effect on the viability of lymphocytes and granulocytes. Exposure to SsESPs significantly increased the splenocyte respiratory burst in response to stimulation with phorbol 12-myristate 13-acetate (PMA), but did not alter baseline ROS production, suggesting that SsESPs enhance splenocyte responsiveness. These findings highlight the complexity of parasite mediated immune changes and indicate that SsESPs have tissue-specific immunomodulatory effects.

## 2. Methods

### 2.1 Breeding and husbandry of wild-caught stickleback

Stickleback were bred from wild-caught fish from Roberts Lake, Gosling Lake, and Sayward Estuary on Vancouver Island, B.C., Canada (Permits: XR422002, NA22-679623) and Kenai Estuary in Alaska (Permits: SF2002-043). Crosses were done in the field and fertilized embryos were transported back to the lab at the University of Massachusetts Lowell. Fish from the same family were housed together at 17ºC in a continuous flow aquarium, adults were stocked at 1 fish/L. These experiments were approved by the University of Massachusetts Lowell IACUC, protocols 18-11-04-Ste and 21-10-07-Ste.

### 2.2 S. solidus collection, breeding, and egg harvest

*S. solidus* were collected from infected stickleback from Gosling, Echo, Boot, and Nimpkish Lakes on Vancouver Island, B. C., Canada (XR2022023, NA23-87881). Infected fish were brought back to the University of Massachusetts Lowell and parasites were dissected and bred to obtain eggs as described in Weber^27^. Eggs were collected on days 1, 3, and 5, washed three times with autoclaved ddH_2_O, and stored in 15 ml of autoclaved ddH_2_O in conical tubes covered in foil at 4ºC.

**Table 1.** Lake Locations.

| Location | Coordinates |
| --- | --- |
| Boot Lake | 50.054056, -125.531981 |
| Echo Lake | 49.986052, -125.410283 |
| Gosling Lake | 50.056825, -125.501542 |
| Nimpkish Lake | 50.512606, -127.016168 |
| Roberts | 50.221132, -125.542559 |
| Sayward Estuary | 50.378552, -125.947912 |
| Kenai River Estuary | 60.541976, -151.219275 |

### 2.3 Experimental Infection

*S. solidus* egg pellet was gently disrupted by inversion, and 200 µl of egg mixture was added to each well of a sterile 6-well plate and filled to the top with autoclaved distilled water. The plate was wrapped in foil and incubated at 20ºC for 3 weeks. The foil was removed and the eggs were exposed to 12 hours of light to induce hatching. Live procercoids were collected and immediately fed to juvenile cyclopoid copepods. 3 weeks post-exposure, copepods were briefly anesthetized in carbonated water and visually screened for *S. solidus* infection at 5.5X magnification. Infected copepods (2-3 per fish) were then fed to lab-raised stickleback originating from Sayward Estuary on Vancouver Island, BC.

### 2.4 Worm and excretory-secretory product collection

Infected fish were euthanized 9-11 weeks post exposure to *S. solidus* infected copepod. To ensure sterility, euthanized fish were disinfected with 70% ethanol in distilled water and dissections and ESP collections were conducted in a biosafety cabinet. Disinfected fish were dissected and *S. solidus* plerocercoids were removed from the body cavity and placed individually in 2 ml of sterile 0.9X PBS. The plerocercoids were then rinsed in fresh, sterile 0.9X PBS for approximately 15 minutes, followed by two additional quick washes in fresh, sterile 0.9X PBS. ESPs were collected by placing cestodes in a sterile, low protein binding 5 ml conical tube (Argos Technologies, T2076S-CA, Jet Bio-Filtration, CFT-002-050, MTC Bio, C2520) filled with 2 ml fresh, sterile 0.9X PBS and incubated at 17ºC for 24 hours. The supernatant containing ESPs was sterile filtered individually using a 0.22 µm PES sterile filter (Nalgene, 725-2520, Fisherbrand, 09-720-511) and the filtered ESPs were stored in low protein binding tubes at -20ºC. After ESP collection each plerocercoid was weighed, and only ESPs of those >50 mg were used in subsequent experiments.

### 2.5 Excretory-secretory sterility testing and concentration

ESPs were individually tested for the presence of gram-positive and -negative bacteria, and endotoxins. Samples of ESPs were cultured on both LB agar (LB agar: Fisher BioReagents™, BP1425-500, petri dishes: Falcon, 351029) and sheep’s blood agar (Remel, R01202) at 17ºC for 24 hours and assessed bacterial content. The Pierce™ Chromogenic Endotoxin Quant Kit (Thermo Scientific™, A39553) was used according to the manufacturer’s instructions to detect low levels of endotoxin (0.01-0.1 EU/mL). Only samples that exhibited no bacterial growth and contained less than 0.01 EU/mL endotoxin were used for downstream analysis.

ESPs of *S. solidus* plerocercoids from fish from the same lake population were pooled together, concentrated, and the final protein concentration was measured. ESPs were placed in the upper chamber of a centrifugal filter unit with a MWCO of 1 kD (Pall Corporation, MCP001C41) and centrifuged for 90 minutes at 3000 g at 4ºC. The final concentration of protein in the pooled sample was measured using a Qubit™ Fluorometer and Qubit™ protein assay (invitrogen Qubit™ Protein Assay Kit, Q33211).

### 2.6 Splenocyte isolation and culture

2-2.5 years old, lab-reared fish were euthanized by an overdose of MS-222 (500 mg/L in ddH_2_O, pH 7.4) for at least 5 minutes, followed by pithing. To reduce contamination, fish were disinfected with 70% ethanol and dissected in the biosafety cabinet. Spleens were pooled and disrupted on a 40 µm filter (Greiner Bio-One, 542140) using a pipet tip and resuspended in fresh, chilled splenocyte culture media (Leibovitz’s L-15 (Medium Gibco, 11415-064), 5% fetal bovine serum (Gibco, 26140079), 2% penicillin-streptomycin (Gibco, 15140122)). Cells were centrifuged for 12 minutes at 90 g and 4ºC. The supernatant was carefully removed via vacuum, pellet was resuspended, the volume measured, and cell concentration and viability was determined using trypan blue exclusion (10% trypan blue in PBS) and a hemocytometer.

### 2.7 Splenocyte viability assessment

Pooled, Roberts Lake splenocytes were plated at a concentration of 1×10^6^ cells/ml in 200 µl per well of a sterile, 96-well tissue culture treated plate (Costar, 3599). Treatment wells received 5, 10, or 20 µg of ESPs, while the control wells received an equivalent volume of diluent (0.9X PBS). ESPs from Gosling, Echo, Boot, and Nimpkish Lakes were used for the viability assays. Heat-denatured ESPs were generated by incubation on a heating block at 95ºC for 20 minutes and cooling for 30 minutes prior to use. The cells and ESPs were incubated for 24-72 hours at 17ºC. Prior to analysis, the plates were placed on ice for 10 minutes to dislodge adherent cells and the cell suspensions were transferred into 1.5 ml microcentrifuge tubes. Cells were spun at 400 g, at 4ºC, for 10 minutes, the supernatant was removed, and the pellet volume was measured. An equivalent volume of DAPI (Sigma-Aldrich, D9542-1MG) diluted 1:1000 in flow cytometry media (0.9X RPMI (Gibco, 11835-030 diluted in cell culture grade water), 3% fetal bovine serum (Gibco, 26140079)) was added to each tube immediately prior to running on an imaging flow cytometer (Cytek Amnis FlowSight).

Flow cytometry data was analyzed in FlowJo™ (v.10.10)^28^. Single cells were gated based on Area and Aspect Ratio. For analysis of whole splenocytes, viable singlets were gated based on DAPI exclusion (Figure 1). For subset analysis, total singlets were separated into subpopulations of lymphocytes, granulocytes, and erythrocytes based on SSC vs Area^29^ and gated based on DAPI exclusion (Figure 1).

**Figure 1.**
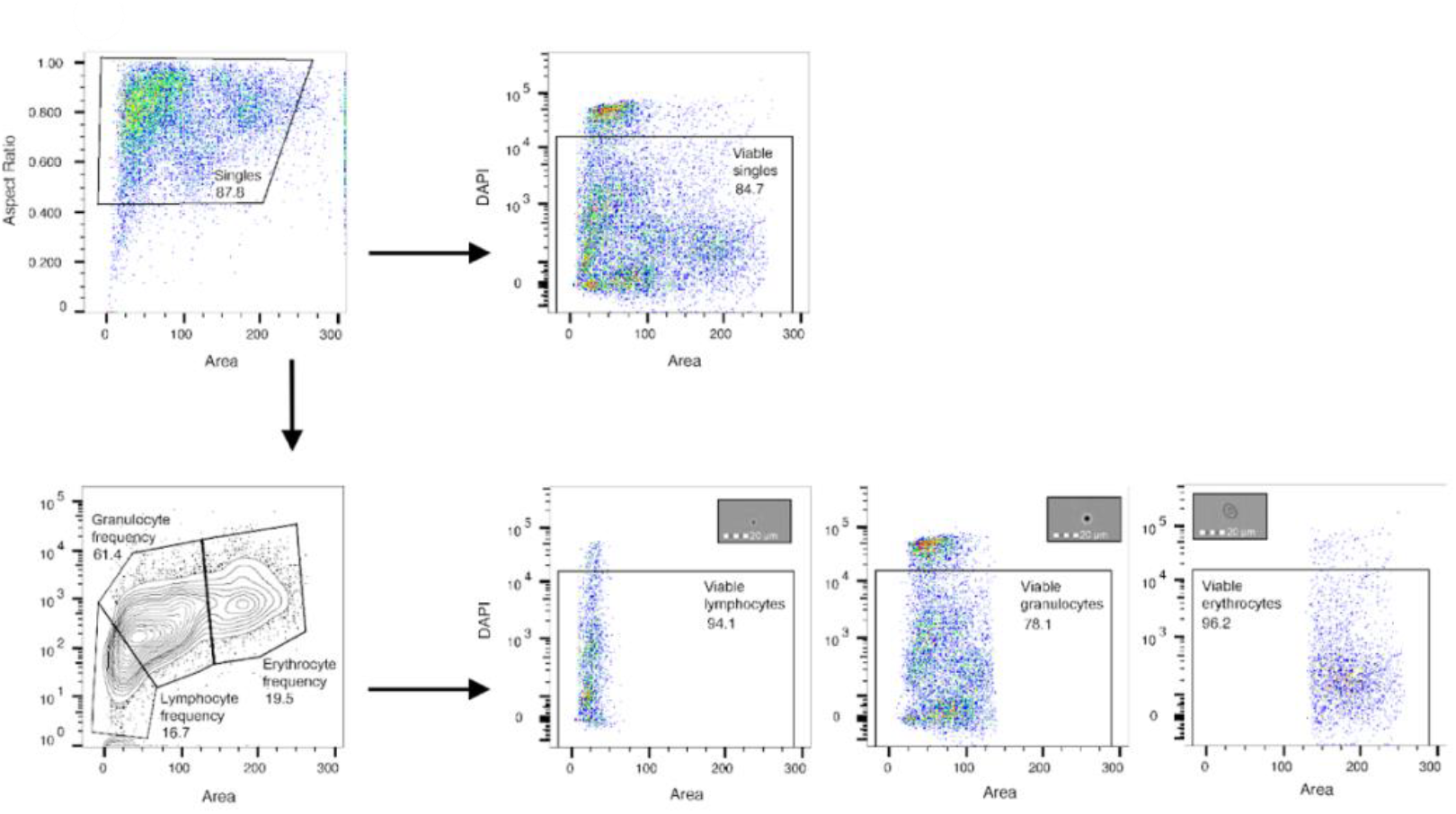
Flow cytometry gating strategy. Single cells were gated based on area and aspect ratio. Viable total splenocyte and subsetted leukocyte populations were gated based on DAPI exclusion. Subpopulations of live singlets were gated based on their SSC (log) and Area (linear) profiles. Leukocyte subsetting strategy was confirmed in a separate experiment using a Cytek Amnis Flowsight imaging flow cytometer (inset images).

### 2.8 ROS assay with stickleback splenocytes

Splenocytes from the Kenai River Estuary fish were plated at a concentration of 5×10^6^ cells/ml in 200 µl per well in a black, clear-bottom 96-well plate (Corning, 3603). Dihydrorhodamine-123 (DHR-123, Sigma-Aldrich, D1054), which oxidizes to rhodamine-123 in the presence of ROS, was used to detect superoxide and hydrogen peroxide. Phorbol 12-myristate 13-acetate (PMA, Stemcell Technologies, 74042), a potent NADPH oxidase activator, was used to stimulate cellular production of ROS. The DHR-123 (0.2 µg/µl final concentration), PMA (0.15 µg/µl final concentration), and 10 µg of ES products (or 0.9X PBS diluent only for control samples) were added to the appropriate wells and incubated at RT for 4 hours. ESP from Echo, Boot, and Nimpkish Lake worms were tested in the ROS assays. Prior to quantification, all wells were mixed thoroughly and fluorescence readings were taken using the Tecan Infinite® F Nano+ microplate reader. Fluorescence was measured at two minute intervals using an excitation/emission laser wavelength at 480/535 nm. Readings were taken from the bottom of the well plate with 10 flashes per reading taken in a filled circular pattern. The gain was set to “optimal”, at 20% relative fluorescent units (RFU).

Readings were taken for 18 minutes to quantify ROS production and the area under the curve (AUC) was calculated. The AUC was calculated using the trapezoidal rule, which divides each curve into segments, finds the area of each individual segment as a trapezoid, and takes the sum of those areas. To find the area of each trapezoidal segment, the following formula was used:

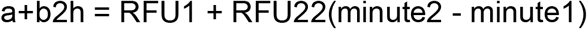

The sum of all segments was found for each curve to obtain the AUC for each sample. Three biological replicates were used for each treatment and the AUC for each replicate were averaged together.

### 2.9 Statistical analysis

Statistical analyses were performed in RStudio (R (4.3.2), RStudio (2023.06.1))^30^. To assess the significant differences in whole splenocyte viability, cell subset viability, and granulocyte/lymphocyte (G/L) ratio between splenocytes exposed to each treatment over time and concentration, two-way ANOVAs were run (‘stats’ package in R). If the ANOVA results showed significant differences, Tukey’s Honestly Significant Difference (HSD) test was run to identify differences in each group. To assess the differences in ROS production, the mean AUC between each treatment group was compared with an unpaired, two-sample t-Test (‘stats’ package in R).

## 3. Results

### 3.1 Canadian *S*. *solidus* ESPs promote stickleback total splenocyte viability

To determine if SsESP-mediated immune modulation extends beyond HKLs and affects other stickleback immune tissues, we measured the viability of stickleback splenocytes cultured with a range of SsESP concentrations (5 µg, 10 µg, and 20 µg) for 24 hours. Viability of splenocytes cultured with SsESPs was significantly higher compared to PBS-only control (two-way ANOVA, F_(1, 28)_ = 6.1, *p* = 0.019, Figure 2A, B). There was no significant effect of concentration (two-way ANOVA, F_(2, 28)_ = 1.77, *p* = 0.189) or interaction between treatment and concentration (two-way ANOVA, F_(2, 28)_ = 0.019, *p* = 0.98).

**Figure 2.**
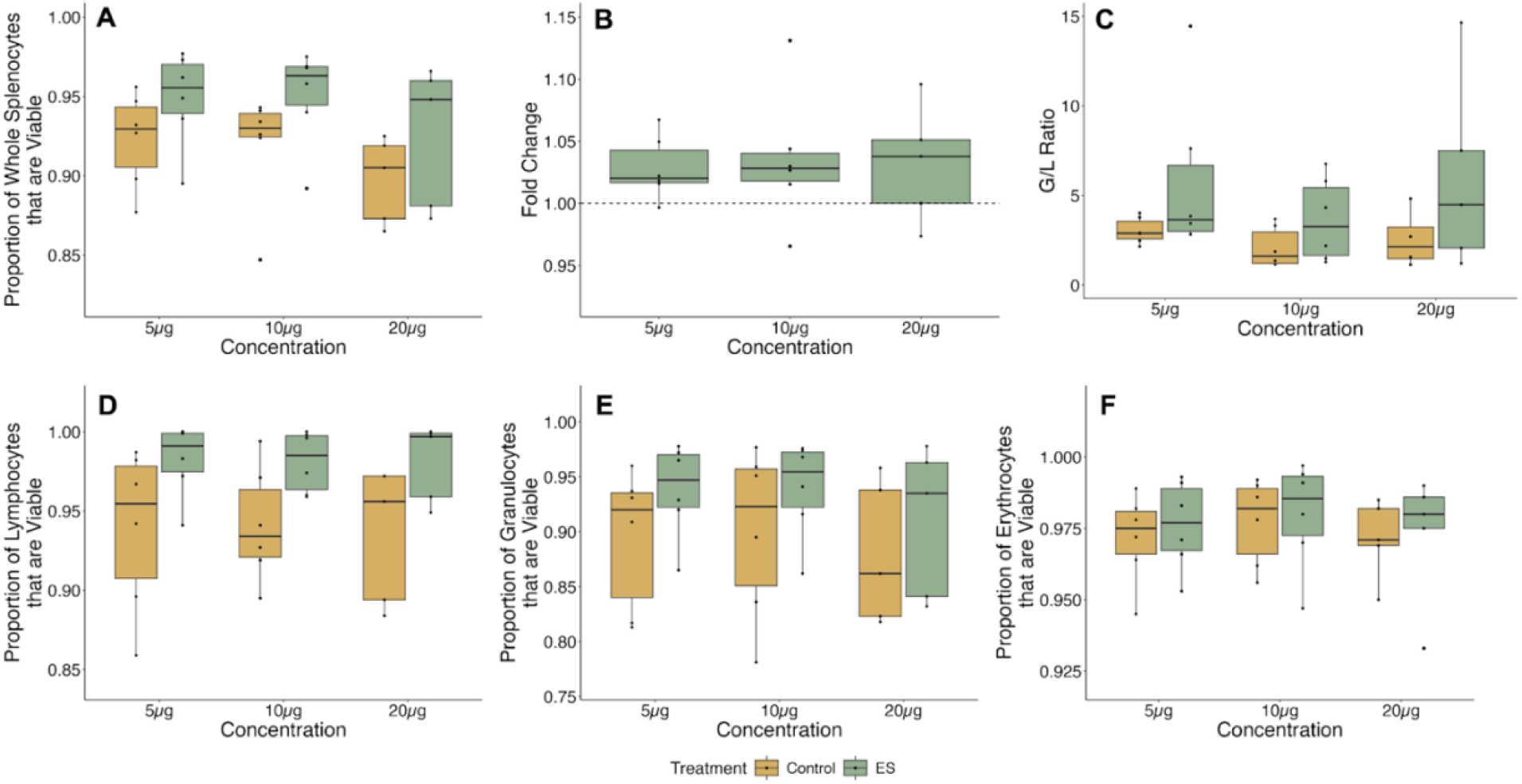
*S. solidus* ESPs increase splenocyte viability and G/L ratio. Proportion of whole splenocytes that are viable (A), fold change in viability (B), and G/L ratio (C) after exposure to 5 µg, 10 µg, and 20 µg of Gosling Lake SsESPs (green boxes) or PBS-only control (yellow boxes). The proportion of viable lymphocytes (D), granulocytes (E), and erythrocytes (F) are also shown. Each dot represents one individual (5 µg (*N* = 6), 10 µg (*N* = 6), and 20 µg (*N* = 5)).

### 3.2 *S. solidus* ESPs promote lymphocyte viability

To assess the immune cell population-specific effects of SsESPs we analyzed the granulocyte/lymphocyte (G/L) ratios of the SsESP-exposed splenocyte cultures. Exposure to ESPs significantly increased the G/L ratio (two-way ANOVA, F_(1, 27)_ = 5.3, *p* = 0.028), even at the lowest concentration (Figure 2C). There was no significant effect of concentration (F_(2, 27)_ = 0.893, *p* = 0.421) or interaction between treatment and concentration (F_(2, 27)_ = 0.247, *p* = 0.783). To assess SsESPs effect on specific immune cell populations, we performed a subset analysis of lymphocytes, granulocytes, and erythrocytes via flow cytometry. Culturing with ESPs significantly increased lymphocyte viability (two-way ANOVA, F_(1, 28)_ = 12.75, *p* = 0.0013, Figure 2D), and did not affect granulocyte (F_(1, 28)_ = 3.30, *p* = 0.080, Figure 2E) and erythrocyte viability (F_(1, 28)_ = 0.24, *p* = 0.628, Figure 2F). For all cell populations, cell visibility was equivalent across SsESP concentrations (lymphocytes; F_(2, 28)_ = 0.022, *p* = 0.979, granulocytes; F_(2, 28)_ = 0.513, *p* = 0.604, erythrocytes; F_(2, 28)_ = 0.428, *p* = 0.656), and the interaction between treatment and concentration was not significant (lymphocytes; F_(2, 28)_ = 0.016, *p* = 0.98, granulocytes; F_(2, 28)_ = 0.035, *p* = 0.966, erythrocytes; F_(2, 28)_ = 0.023, *p* = 0.977).

### 3.3 SsESP-mediated lymphocyte viability enhancement is more pronounced over time and is not diminished with heat denaturation

To determine if the viability-enhancing effects of SsESPs are sustained over time we incubated whole splenocytes with 10 µg of SsESP protein for 24, 48, and 72 hours. To determine if viability enhancement was mediated by protein components of the excretory secretory products, we also exposed splenocytes to 10 µg of heat-denatured SsESPs for 24, 48, and 72 hours. Whole splenocyte viability was not significantly affected by treatment (two-way ANOVA, F_(2, 18)_ = 2.88, *p* = 0.08), time (F_(2, 18)_ = 0.50, *p* = 0.62), or their interaction (F_(4, 18)_ = 0.12, *p* = 0.98) (Figure 3A, B). However, treatment had a significant effect on the G/L ratio (two-way ANOVA, F_(2, 17)_ = 8.06, *p* = 0.0035) (Figure 3C). Splenocytes cultured with SsESPs had a higher G/L ratio than those cultured with control media (Tukey’s, *p* = 0.015) or heat-denatured SsESPs (*p* = 0.005). The G/L ratio did not change over time (F_(2, 17)_ = 0.91, *p* = 0.42) and the interaction between treatment and timepoint (F_(4, 17)_ = 0.87, *p* = 0.50) did not significantly affect the G/L ratio (Figure 3C).

**Figure 3.**
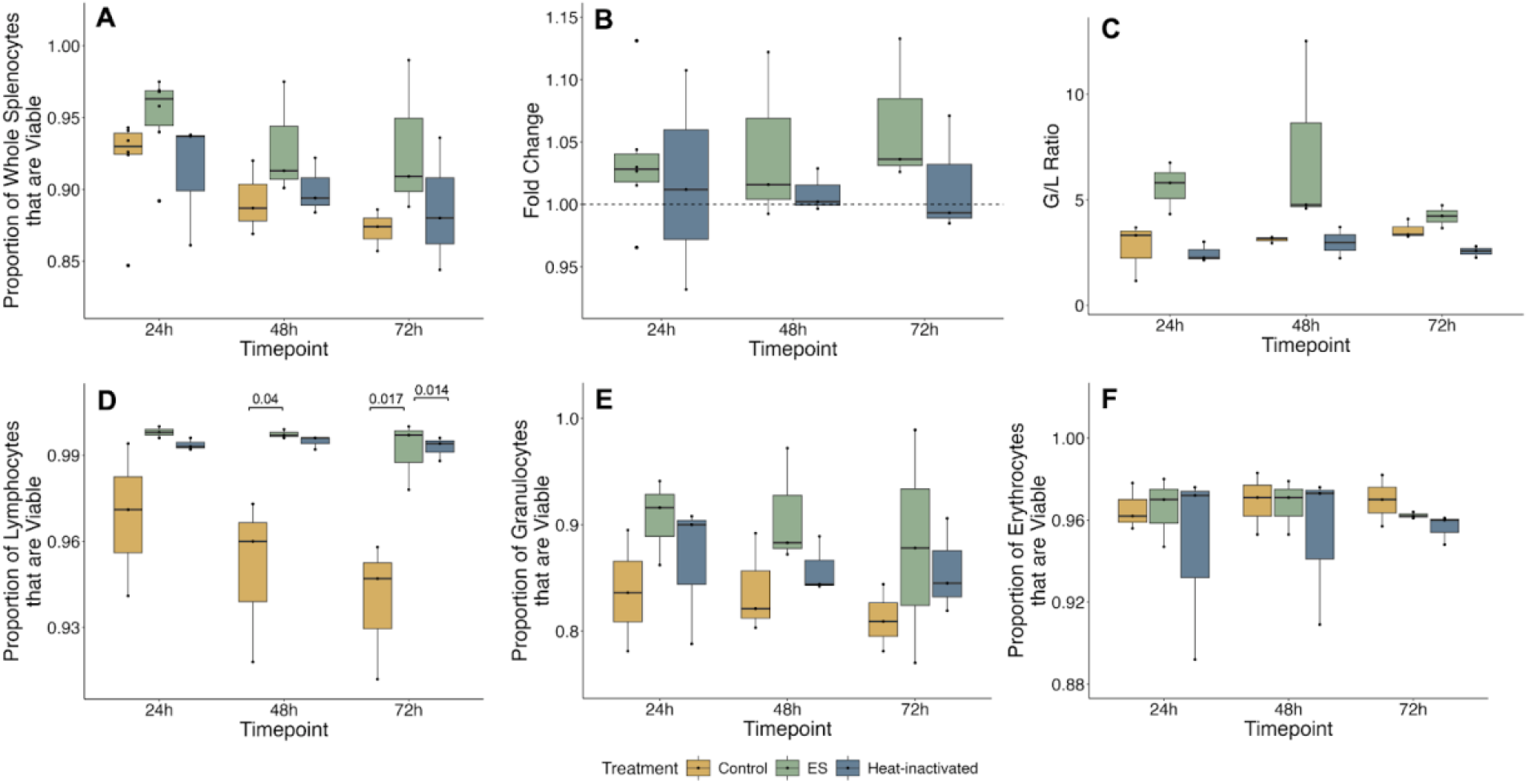
*S. solidus* ESPs increase lymphocyte viability for 72 hours in culture. Proportion of whole splenocytes that are viable (A), fold change in viability (B), and G/L ratio (C) after exposure to Gosling Lake SsESPs (green boxes), heat-inactivated SsESPs (blue boxes), or PBS-only control (yellow boxes) for 24, 48, and 72 hours (A). The proportion of viable lymphocytes (D), granulocytes (E), and erythrocytes (F) are also shown. Significant changes in lymphocyte viability with exposure to SsESPs and heat-inactivated SsESPs are indicated by p-values (D). Each dot represents one individual (A-B; 24h (*N* = 6), 48h (*N* = 3), and 72h (*N* = 3), C-F; 24h (*N* = 3), 48h (*N* = 3), and 72h (*N* = 3)).

Treatment with SsESPs significantly enhanced the viability of splenic lymphocytes over long-term exposure (two-way ANOVA, F_(2, 18)_ = 20.88, *p* = 2.04e-05) (Figure 3D). Treatment with both SsESPs (Tukey’s, *p* = 5.66e-05) and heat-denatured ESPs (*p* = 9.87e-05) led to similar increases in lymphocyte viability (Figure 3D). SsESP-induced lymphocyte viability was highest at 48 (Tukey’s, *p* = 0.04) and 72 hours (*p* = 0.017) (Figure 3D). At 72 hours, the viability of lymphocytes exposure to heat-denatured ESPs was also significantly greater than those exposed to control media (*p* = 0.014) (Figure 3D). Timepoint (F_(2, 18)_ = 1.35, *p* = 0.29) and the interaction between treatment and timepoint (F_(4, 18)_ = 0.73, *p* = 0.59) did not significantly affect viability (Figure 3D).

Granulocyte viability increased with exposure to SsESPs but was not overall affected by treatment (two-way ANOVA, F_(2, 18)_ = 3.19, *p* = 0.065), timepoint (F_(2, 18)_ = 0.36, *p* = 0.7), or their interaction (F_(4, 18)_ = 0.06, *p* = 0.99) (Figure 3E). Erythrocyte viability was not affected by treatment (two-way ANOVA, F_(2, 18)_ = 1.28, *p* = 0.3), timepoint (F_(2, 18)_ = 0.08, *p* = 0.924), or their interaction (F_(4, 18)_ = 0.07, *p* = 0.99) (Figure 3F).

### 3.4 SsESPs from four Canadian populations demonstrate equivalent enhancement of splenocyte viability

To determine population-level variation in SsESPs, we compared ESPs from *S. solidus* from four Canadian lakes (Gosling, Echo, Boot, and Nimpkish). Stickleback splenocytes were incubated in the presence or absence of 20 µg of SsESPs for 24 hours and their viability was assessed via flow cytometry, and the fold change in viability was determined. Similar fold changes in whole splenocyte viability were observed with exposure to SsESPs from all four lakes (Figure 4A). A positive fold change in the G/L ratio was observed with exposure to Gosling and Nimpkish Lake SsESPs, while exposure to Boot and Echo Lake SsESPs resulted in a negative fold change in G/L ratio (Figure 4B). Similar positive fold changes of lymphocytes (Figure 4C) and granulocytes (Figure 4D) that are viable, and neutral fold changes in erythrocytes (Figure 4E) that are viable were found with exposure to SsESPs from all lakes.

**Figure 4.**
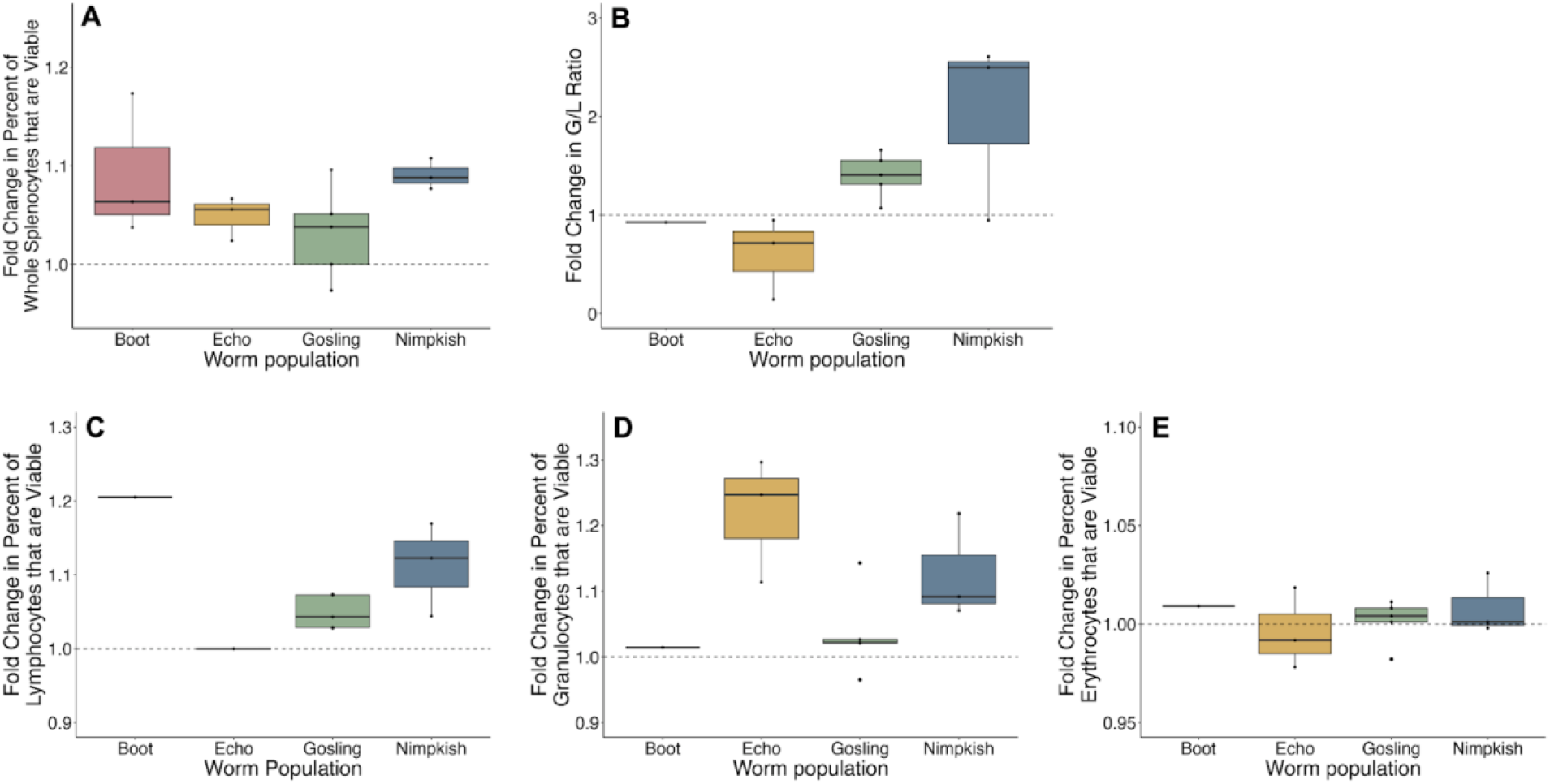
*S. solidus* ESPs from Canadian lakes induce similar fold changes in viability but target different cell types. Fold change in percent of whole splenocytes that are viable after exposure to 20 µg SsESPs from Boot Lake (pink, *N* = 3), Echo Lake (yellow, *N* = 3), Gosling Lake (green, *N* = 5), and Nimpkish Lake (blue, *N* = 3) (A). Fold change in G/L ratio after exposure to 20 µg SsESPs from Boot Lake (*N* = 1), Echo Lake (*N* = 3), Gosling Lake (*N* = 5), and Nimpkish Lake (*N* = 3) (B). Fold change in percent of viable lymphocytes (C), granulocytes (D), and erythrocytes (E) after exposure to Boot Lake (*N* = 1), Echo Lake (*N* = 3), Gosling Lake (*N* = 5), and Nimpkish Lake (*N* = 3) SsESPs.

### 3.5 ROS increase in stickleback splenocytes

To assess *S. solidus*’s effect on stickleback splenocyte function, we measured the ROS production of splenocytes cultured in the presence or absence of SsESPs from three Canadian populations: Echo, Boot and Nimpkish Lakes. Cells that received SsESPs alone without PMA stimulation produced minimal amounts of ROS (AUC 1193.96 ± 119.6 (mean ± SE)) equivalent to unstimulated controls (1011.98 ± 51.9) (*p* = 0.15) (Figure 5). PMA stimulation with exposure to SsESPs from Echo (Welch’s t-test, *p* = 0.0005), Boot (*p* = 0.002), and Nimpkish (*p* = 0.0002) Lakes caused greater ROS production than stimulation with PMA alone (Figure 5). ROS production in these cells remained elevated for the entire duration of the experiment.

**Figure 5.**
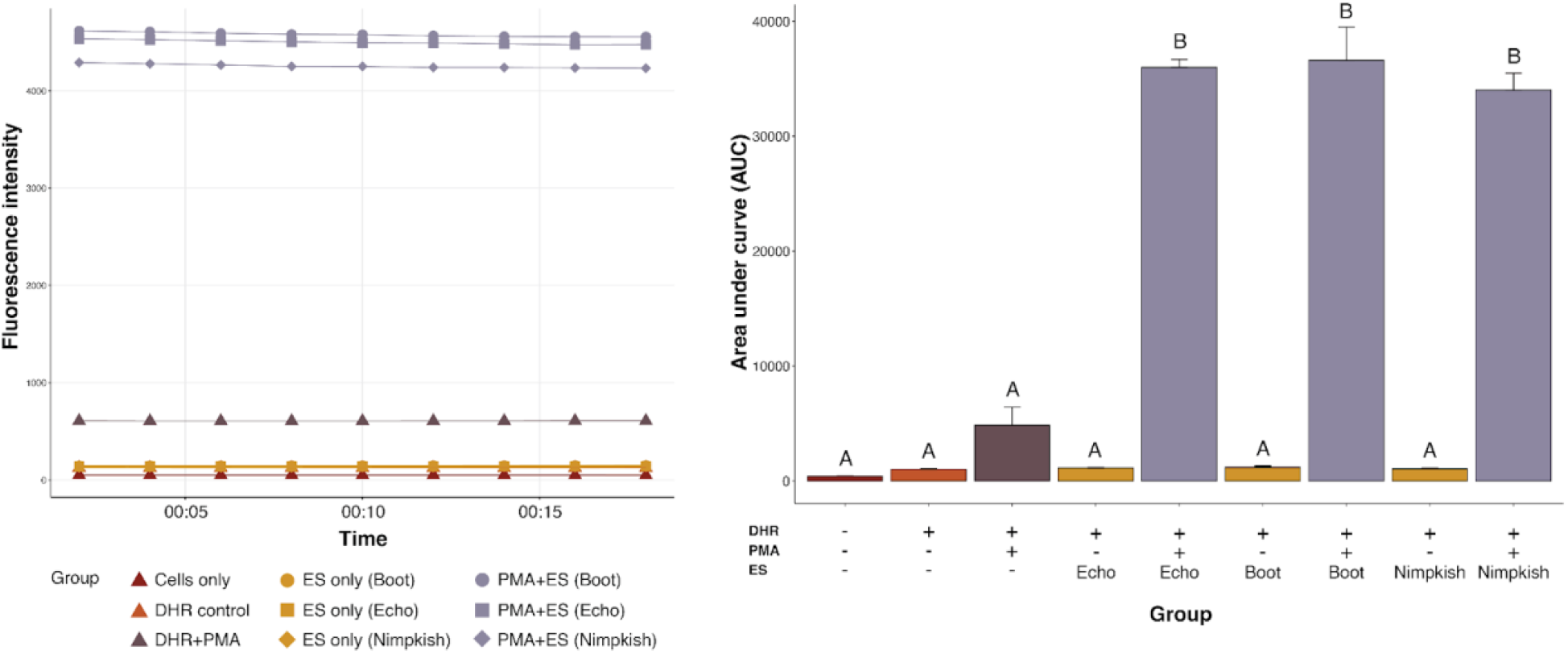
SsESPs increase splenocyte ROS production following PMA stimulation. The area under the curve (AUC) of the fluorescence intensity of stickleback splenocytes stimulated with SsESPs from Echo, Boot, and Nimpkish Lakes. Significant differences between treatments are noted with different letters. Splenocytes from the same individual received each treatment (*N* = 3).

## 4. Discussion

To expand our understanding of *S. solidus* influence on host immunity, we assessed threespine stickleback splenocyte viability and function when cultured with SsESPs from several Canadian lakes. We observed that Canadian SsESPs enhance splenocyte viability, that this effect is detectable at low concentrations and is maintained over 72 hours in culture. We originally hypothesized that *S. solidus* would suppress stickleback immunity by reducing splenocyte viability, as many helminths are known to induce host cell death through specific apoptotic ESP proteins (reviewed in Zakeri, A. 2017^31^). However, inducing host cell death may not increase parasite fitness if doing so also decreases host fitness. *S. solidus* reaches full sexual maturity in the stickleback host, therefore host condition is paramount to *S. solidus*’ reproductive success. Promoting host cell viability is a potential strategy that may reduce host morbidity, allowing for transmission of *S. solidus* to its terminal host.

For these experiments, we analyzed whole splenocyte samples which are composed of many different cell populations. Though tools to assess splenic populations are lacking, our analysis of splenocytes by FSC and SSC revealed that the relative ratio of granulocytes to lymphocytes increased in culture with SsESPs. The observed increase in this ratio could be reflective of A) an expansion of granulocyte populations or B) a reduction of lymphocyte populations. Viability analysis on splenocyte subsets indicated that SsESPs enhance splenic lymphocyte viability and had no effect on granulocytes or erythrocytes viability suggesting the former scenario is more likely. An expansion of granulocytes likely indicates activation or mobilization of the innate immune response, which is generally expected during early exposure to parasites in fish^32^. Further analysis of the granulocyte/lymphocyte populations upon exposure to SsESPs is needed to draw conclusions about whether this is a mechanism of innate immune activation, or an adaptive immunosuppression strategy mediated by *S. solidus*.

Our comparison of cestodes from four Canadian lakes showed equivalent SsESP-induced viability enhancement. This contrasts with a previous report comparing SsESPs from three European *S. solidus* populations that showed an effect of parasite origin on HKL viability^24^. The lack of variation we observed among Canadian SsESPs could be due to the relatively close proximity of the lakes studied. Dispersal of *S solidus* eggs via the terminal bird host may facilitate gene flow between nearby lakes, and *S. solidus* genetic landscape is highly connected to watershed boundaries^33,34^. Gosling, Boot, and Echo Lakes are all in the Campbell river watershed, while Nimpkish Lake is the farthest geographically in Tiakwa Creek Watershed^35^. Therefore, to determine if population-level variation exists in SsESP-induced splenocyte viability enhancement, future work should examine North American cestodes from more distant locales.

The respiratory burst is an essential antimicrobial and antiparasite response primarily mediated by neutrophils and macrophages, which may extend to red blood cells (RBCs) in teleosts^25,36^. Previous study in stickleback HKL cultures showed that SsESPs initially upregulate ROS production, but by 96 hours ROS production is suppressed^24^. This immune-suppressive effect is likely mediated via superoxide dismutase (SOD), peroxidases, and other proteins that may function to reduce oxidative stress that were recently identified in the *S. solidus* secretome^22^. We found that while SsESPs alone did not trigger splenocyte ROS release, however similar to HKL, culturing with SsESPs primed splenocytes to produce high amounts of ROS when stimulated with PMA. This enhanced sensitivity to respiratory burst induction is surprising considering that ROS-induced tissue damage can reduce both host and parasite fitness^37^. However, it is possible that maintaining/enhancing host ROS production could protect the host from co-infection, and therefore this could be an indirect mechanism used by the parasite to increase its own fitness by maintaining the innate immune responses of its host. Unfortunately, the limited nature of the SsESPs prevented us from exploring the effects of SsESPs on splenic ROS production at later time points.

Our viability results in splenocytes contrast with previous study of head kidney leukocytes (HKL). This work showed that, depending on the origin of the SsESPs, either a reduction or no change in cultured HKL viability^24^. There are several potential factors that could contribute to this difference. First, as HK and spleen have different immune cell composition and function, these divergent results suggest that SsESPs may have tissue-specific effects. The HK is primarily a hematopoietic organ which generates leukocytes and has a high frequency of innate immune cells^25^, while the spleen is a secondary lymphoid organ containing mature lymphocytes undergoing activation and clonal expansion^38^. It is possible that SsESPs have differential effects on developing vs mature lymphocytes. Second, differences in viability between HKL and splenocytes cultured with SsESPs could also be explained by differences in ESP preparation and confirmation methods. For the present study, SsESPs were collected sterilely and only those supernatants that passed both an endotoxin screen and culture tests were used in *in vitro* assays. Furthermore, to normalize across experiments, we used a standardized SsESP concentration across all assays. Therefore, these technical modifications could contribute to the differences observed between our splenocyte and previously published HKL experiments. Lastly, differences between our results and previous reports in HKL could reflect population level variation among the cestodes and/or fish populations used. Our study used Canadian cestodes and fish, whereas the HKL experiments used European populations of hosts and parasites^24^. It is possible that one or more of these factors contributed to the differences in SsESP-induced viability changes between spleen and head kidney. Unfortunately, the extremely limited nature of Canadian SsESPs prevented us from exploring these options further.

## Acknowledgements

We thank Trey Sasser from the University of Alaska Anchorage and Dr. Cole Wolf from the University of Wisconsin Madison for their help with stickleback collections in 2022 and 2023 respectively. A special thank you to the budding scientist, Jordan, at Echo Lake who helped us collect wormy stickleback with her toy net in 2023. Thank you also to Jack Lepine and Sonny Farfan at the UML Core Research Facilities for technical support. This project was funded by NIH R01 AI146168-01.

